# Ceragenins and antimicrobial peptides kill bacteria through distinct mechanisms

**DOI:** 10.1101/2020.10.20.346411

**Authors:** Gabriel Mitchell, Melanie R. Silvis, Kelsey C. Talkington, Jonathan M. Budzik, Claire E. Dodd, Justin M. Paluba, Erika A. Oki, Kristine L. Trotta, Daniel J. Licht, David Jimenez-Morales, Seemay Chou, Paul B. Savage, Carol A. Gross, Michael A. Marletta, Jeffery S. Cox

## Abstract

Ceragenins are a family of synthetic amphipathic molecules designed to mimic the properties of naturally-occurring cationic antimicrobial peptides (CAMPs). Although ceragenins have potent antimicrobial activity, whether their mode of action is similar to that of CAMPs has remained elusive. Here we report the results of a comparative study of the bacterial responses to two well-studied CAMPs, LL37 and colistin, and two ceragenins with related structures, CSA13 and CSA131. Using transcriptomic and proteomic analyses, we found that *Escherichia coli* responds similarly to both CAMPs and ceragenins by inducing a Cpx envelope stress response. However, whereas *E. coli* exposed to CAMPs increased expression of genes involved in colanic acid biosynthesis, bacteria exposed to ceragenins specifically modulated functions related to phosphate transport, indicating distinct mechanisms of action between these two classes of molecules. Although traditional genetic approaches failed to identify genes that confer high-level resistance to ceragenins, using a Clustered Regularly Interspaced Short Palindromic Repeats interference (CRISPRi) approach we identified *E. coli* essential genes that when knocked down modify sensitivity to these molecules. Comparison of the essential gene-antibiotic interactions for each of the CAMPs and ceragenins identified both overlapping and distinct dependencies for their antimicrobial activities. Overall, this study indicates that while some bacterial responses to ceragenins overlap with those induced by naturally-occurring CAMPs, these synthetic molecules target the bacterial envelope using a distinctive mode of action.

**IMPORTANCE:** The development of novel antibiotics is essential since the current arsenal of antimicrobials will soon be ineffective due to the widespread occurrence of antibiotic resistance. Development of naturally-occurring cationic antimicrobial peptides (CAMPs) for therapeutics to combat antibiotic resistance has been hampered by high production costs and protease sensitivity, among other factors. The ceragenins are a family of synthetic CAMP mimics that kill a broad spectrum of bacterial species but are less expensive to produce, resistant to proteolytic degradation and have been associated with low levels of resistance. Determining how ceragenins function may identify new essential biological pathways of bacteria that are less prone to development of resistance and will further our understanding of the design principles for maximizing the effects of synthetic CAMPs.

## INTRODUCTION

Our current arsenal of antibiotics will soon be ineffective against the simplest bacterial infections due to the continued spread of antibiotic resistance (AR) (1). AR has been identified in virtually all bacterial species of clinical relevance, including Gram-positive and Gram-negative bacteria as well as mycobacteria (2). Despite the threat that AR represents to global health, there is a lack in the development of antimicrobials with innovative mechanisms of action (3–5). A better understanding of the fundamental principles of how antibiotics kill microbes and how AR develops will help break the futile cycle of antibiotic development and microbial evolution.

Antimicrobial peptides are structurally diverse molecules expressed in a wide array of organisms that directly kill microbes, including bacteria (6, 7). Many antimicrobial peptides, such as the class of cationic antimicrobial peptides (CAMP), rapidly kill bacteria by disrupting membranes although other mechanisms of action were also suggested (7–9). The potential of using CAMPs to treat AR infections has become a research focus due to their action against a broad spectrum of pathogens, their selectivity toward microbial membranes and the low appearance of resistance (6, 10). Despite some progress in this area, significant barriers to CAMP therapeutic development include high production costs, toxicity, susceptibility to proteolytic degradation and activation of allergic responses (6, 10).

Ceragenins are a family of synthetic amphipathic molecules derived from cholic acid designed to mimic the activity of endogenous CAMPs (11, 12). These molecules are inexpensive to manufacture and are not susceptible to proteolysis, making them an attractive alternative to peptide-based synthetic CAMPs. Importantly, ceragenins have antimicrobial activity against a broad spectrum of microbes, which include both Gram-negative and Gram-positive bacteria (11, 13). High-level resistance to ceragenins is seemingly difficult to acquire in the lab as attempts to isolate ceragenin-resistance bacterial mutants failed in the Gram-positive bacterium *Staphylococcus aureus* and identified only modest and unstable resistance in Gram-negatives (14). Although ceragenins were designed as CAMP mimics and can depolarize bacterial membranes (15), the inability to identify bona-fide ceragenin-resistant bacterial mutants represents a major barrier in understanding their mechanism of action.

Here we take a comparative approach using a combination of transcriptomic, proteomic and genetic approaches to compare the bacterial responses to treatment with ceragenins and two well-studied CAMPs. The results of this study suggests that ceragenins kill bacteria by disrupting the bacterial envelope through a distinctive mode of action from naturally-occurring CAMPs. We also show, for the first time, that ceragenins have activity against mycobacteria despite their distinctive cell wall architecture.

## MATERIALS AND METHODS

### Antimicrobial compounds

CSA13 and CSA131 (16) as well as CSA44 and CSA144 (17) were prepared as described previously and solubilized at 10 mg/mL in sterile distilled and deionized (DD) water. LL37 (Anaspec, Fremont, California, USA), colistin (Sigma-Aldrich, St-Louis, MO, USA) and ciprofloxacin (MP Biomedicals, Irvine, California, USA) were solubilized at 10 mg/mL in sterile DD water. Erythromycin (Sigma-Aldrich) was solubilized at 10mg/mL in ethanol. Antimicrobial compounds were aliquoted and stored at −20° C. Freeze-thaw cycles of stock solutions were limited to three times.

### Bacterial strains, growth conditions

*E. coli* MG1655 (18), *L. monocytogenes* 10403S (19), *M. marinum* strain M (20), *M. smegmatis* mc^2^ 155 (21) and *M. tuberculosis* Erdman (20) were as previously described. *M. avium* mc^2^ 2500 is a clinical strain isolated from an acquired immunodeficiency syndrome (AIDS) patient with pulmonary disease and predominantly formed a smooth/transparent colony morphotype on solid agar (22). *M. avium* mc^2^ 2500D is an isogenic, laboratory-derived strain with an opaque colony morphotype. *E. coli* and *L. monocytogenes* were routinely grown on Mueller-Hinton agar (MHA) (BD, Franklin Lakes, New Jersey, USA) plates or in cation-adjusted Mueller-Hinton (CAMH) (BD) broth and on Brain-heart infusion agar (BHIA) (Becton Dickinson) plates or in Brain-heart infusion (BHI) (BD) broth, respectively. *M. avium* and *M. tuberculosis* were cultured in Middlebrook 7H9 (BD) broth containing 0.5% glycerol, 10 % Oleic Albumin Dextrose Catalase (OADC) (Sigma-Aldrich) and 0.05% Tween-80. *M. marium* was cultured in Middlebrook 7H9 broth containing 0.5% glycerol, 10 % OADC and 0.2% Tween-80. *M. smegmatis* was grown on Middlebrook 7H10 (BD) plates or Middlebrook 7H9 plates containing 0.5% glycerol, 0.5% dextrose and 0.2% Tween-80, unless otherwise stated. All bacterial strains were grown at 37° C, except from *M. marinum*, which was grown at 30° C. Liquid cultures were incubated with shaking, unless otherwise stated.

### Antibiotic susceptibility testing

Minimal inhibitory concentrations (MICs) of antimicrobial compounds against *E. coli* and *L. monocytogenes* were determined by a broth microdilution technique following the recommendations of the Clinical and Laboratory Standards Institute (CLSI) (23), except that BHI was used to perform assays on *L. monocytogenes*. Antibiotic quality control experiments were performed using *E. coli* ATCC25922 (ATCC, Manassas, VA, USA). A similar protocol using extended incubation periods was used to determine MICs against *M. avium* (10 days), *M. marinum* (5 days), *M. smegmatis* (3 days) and *M. tuberculosis* (14 days). For the determination of MICs using mycobacterial species, plates were placed in a vented container containing damp wipes to minimize evaporation.

### Time-kill experiments

Time-kill experiments were performed to characterize the effect of compounds on bacterial growth and survival. Bacteria were inoculated at 10^5^-10^6^ CFU/mL in liquid media in the absence or presence of antibiotics at the following concentrations: *E. coli*, 0.5 μg/mL colistin, 64 μg/mL LL37, 4 μg/mL CSA13 and 4 μg/mL CSA131; *L. monocytogenes*, 2 μg/mL CSA13, 2 μg/mL CSA131, 2 μg/mL ciprofloxacin and 0.25 μg/mL erythromycin; *M. smegmatis*, 0.5 μg/mL CSA13 and 0.5 μg/mL ciprofloxacin. Bacterial cultures were grown at 37° C with shaking and the number of CFU/mL was determined at several time points. Plates without Tween-80 were used for the CFU determination of *M. smegmatis* cultures.

### Serial passage experiments

Serial passage of bacteria in presence of sub-inhibitory concentrations was performed as previously described (14, 24). Experiments were performed in CAMH or BHI broth for *E. coli* and *L. monocytogenes*, respectively. In few cases, bacteria growing at the two highest sub-inhibitory concentrations of antimicrobial had to be combined in order to get an inoculums of 10^5^-10^6^ CFU/mL.

### Preparation and sampling of bacterial cultures for transcriptomic profiling

A single colony of *E. coli* MG1655 was inoculated into CAMH broth and incubate 16-18 h at 37° C with shaking. Cultures were diluted in fresh media to an *A_600nm_* of 0.1, incubated at 37° C with shaking until an *A_600nm_* of 0.8-1.0 (~2h) and antibiotics were added to each cultures, which were further incubated for 1 h at 37° C with shaking. Antibiotic were adjusted to concentrations having similar impact on *E. coli* growth for that particular, higher bacterial density, culture format (i.e. 4 μg/mL colistin, 8 μg/mL CSA13, 8 μg/mL CSA131 and 256 μg/mL LL37). Cultures samples were then mixed 1:2 with RNA Protect Bacteria Reagent (QIAGEN, Germantown, Maryland, USA), vortexed immediately for 5 seconds and incubated for 5 min at room temperature. The bacterial suspensions were centrifugated for 10 min at 5,000 × g, supernatants discarded and pellets were stored few days at −80° C before proceeding to RNA extraction.

### RNA purification and sequencing

Bacterial pellets were resuspended in 100 μL of 10 mM Tris, 1 mM EDTA, pH 8.0 buffer containing 10 mg/mL lysozyme (Sigma-Aldrich). 2.5 μL of 20 mg/mL Proteinase K (NEB, Ipswich, Massachusetts, USA) was added and samples were incubated at room temperature for 10 min, with frequent mixing. Samples were combined with 0.5 μL of 10% SDS and 350 μL of Lysis Buffer (Ambion life technologies, Invitrogen, Carlsbad, California, USA) containing β-mercaptoethanol, vortexed and the lysate was transferred into a 1.5 mL RNase-free microcentrifuge tube. Samples were then passed 5 times through an 18-21-gauge needle and centrifuged at 12,000 × g for 2 minutes at room temperature. Supernatants were transferred to a new 1.5 mL RNase-free microcentrifuge tube before proceeding to the washing and elution steps described in the PureLink RNA Mini Kit (Ambion life technologies). Samples were treated with DNase (NEB) for 15 min at 37° C in a volume of 50 μL and 5 μL of 25 mM EDTA was added. Samples were further incubated 10 min at 75° C and quickly placed on ice before being cleaned and re-eluted using the RNA Clean & Concentrator kit (Zymo Research, Irvine, CA, USA) and stored at −80° C. The quality and the quantity of each RNA samples was analyzed by the UC Berkeley QB3 facility using a bioanalyzer and the Qubit technology. RNA samples were sequenced and preliminary analyzed by the UC Davis Genome Center and the UC Davis Bioinformatics Core.

### Analysis of RNAseq data

The differential expression analyses were conducted using the limma-voom Bioconductor pipeline (25) (EdgeR version 3.20.9, limma version 3.34.9) and R 3.4.4 by the UC Davis Bioinformatics Core. The multidimensional plot was created using the EdgeR function plotMDS. Pathway analyses were performed using DAVID Bioinformatics Resources 6.8 (26, 27). Only annotation terms from the following databases were included: UP (UniProt) Keywords, COG (Cluster of Orthologous Groups) Ontology, GO (Gene Ontology for Biological process, Molecular function and Cellular component) and KEGG (Kyoto Encyclopedia of Genes and Genomes). Venn diagram analyses were performed using the tool provided on the Bioinformatics & Evolutionary Genomics website of Ghent University (28). Promoter and regulatory binding analyses were performed using the Gene Expression Analysis Tools (29). Only one repeated binding sites and promoters was considered for each gene for any specific transcription factors. Lists of genes to be included in specific regulons were retrieved from the RegulonDB Database (30). Fold changes for few transcripts of regulons CpxR (*cpxQ, csgC, cyaR, efeU, rprA, rseD*), PurR (*codA, codB*) and PhoB (*cusC,phnE,prpR*) were not included in this analysis. The information related to the expected activity (induction and/or repression) of each transcription factor on specific genes were also retrieved from the RegulonDB Database.

### Protein extraction and peptide preparation from *E. coli* cultures

A single colony of *E. coli* MG1655 was inoculated into CAMH broth and incubate 16-18 h at 37° C with shaking. Cultures were diluted in fresh media to an *A_600nm_* of 0.1, incubated at 37° C with shaking until an *A_600nm_* of 0.8-1.0 (~2 h) and antibiotics (4 μg/mL colistin or 8 μg/mL CSA13) were then added to each cultures, which were further incubated for 3 h at 37° C with shaking. Protein were extracted, digested and desalted, as previously described (31), with few modifications. Briefly, 23 mL of bacteria cultures were washed twice in cold PBS and resuspended in 4 mL of lysis buffer (8 M urea, 150 mM NaCl, 100 mM ammonium bicarbonate, pH 8) containing Roche mini-complete protease inhibitor EDTA-free and Roche PhosSTOP (1 tablet of each per 10 mL of buffer) (Roche, Basel, Switzerland). Samples (on ice) were then sonicated 10 times with a Sonics VibraCell probe tip sonicator at 7 watts for 10 seconds. Insoluble precipitates were removed from lysates using a 30 min centrifugation at ~16,100 × g at 4° C and the protein concentration of each lysates was determined using the microplate procedure of the Micro BCA™ Protein Assay Kit (Thermo Fischer Scientific, Emeryville, CA, USA). Clarified lysates (1 mg each) was reduced with 4 mM tris(2-carboxyethyl)phosphine for 30 min at room temperature, alkylated with 10 mM of iodoacetamide for 30 min at room temperature in the dark and quenched with 10 mM 1,4-dithiothreitol for 30 min at room temperature in the dark. Samples were diluted with three volumes of 100 mM ammonium bicarbonate, pH 8.0, and incubated with 10 μg of sequencing grade modified trypsin (Promega, Madison, WI, USA) while rotating at room temperature for 18 hours. Trifluoroacetic acid (TCA) was then added to a final concentration of 0.3% to each samples, followed by 1:100 of 6M HCl and the removal of insoluble material by centrifugation at ~2,000 × g for 10 min. SepPak C18 solid-phase extraction cartridges (Waters, Milford, MA, USA) were activated with 1 mL of 80% acetonitrile (ACN), 0.1% TFA, and equilibrated with 3 mL of 0.1% TFA. Peptides were desalted by applying samples to equilibrated columns, followed by a washing step with 3 mL of 0.1% TFA and elution with 1.1 mL of 40% ACN, 0.1% TFA. The subsequent global protein analysis was performed using 10 μg of each desalted peptide sample.

### Liquid chromatography, mass spectroscopy and label-free quantification

Peptides were analyzed using liquid chromatography and mass spectroscopy, as previously described (31). Mass spectrometry data was assigned to *E. coli* sequences and MS1 intensities were extracted with MaxQuant (version 1.6.0.16) (32). Data were searched against the *E. coli* (strain K12) protein database (downloaded on November 6, 2018). MaxQuant settings were left at the default except that trypsin (KR|P) was selected, allowing for up to two missed cleavages. Data were then further analyzed with the artMS Bioconductor package (33), using the MSstats Bioconductor package (version 3.14.1) (34) and the artMS version 0.9. Contaminants and decoy hits were removed, and samples were normalized across fractions by median-centering the log_2_-transformed MS1 intensity distributions. The MSstats group comparison function was run with no interaction terms for missing values, no interference, unequal intensity feature variance as well as restricted technical and biological scope of replication. Log_2_(fold change) for protein/sites with missing values in one condition but found in > 2 biological replicates of the other condition of any given comparison were estimated by imputing intensity values from the lowest observed MS1-intensity across samples (33), and *P*_values_ were randomly assigned between 0.05 and 0.01 for illustration purposes.

### Identification of genetic determinants of resistance to antibiotics using CRISPRi

A pooled CRISPRi library of strains with inducible knockdown of genes predicted to be essential (FIG. S1) was used to study the genetic determinants of resistance to CAMPs and ceragenins. To quantify the antibiotic sensitivity of each CRISPRi strain, the relative proportion of each sgRNA spacer in the mixed population was enumerated by deep sequencing, after 15 doublings in presence of saturating IPTG and 0.031 μg/mL colistin, 12 μg/mL LL37, 0.5 μg/mL CSA13 and 0.25 μg/mL CSA131. Briefly, a single glycerol stock of the pooled library was fully thawed, inoculated into 10 mL LB at 0.01 *A*_600_, and grown for 2.5 hr (final ~0.3 *A*_600_) at 37° C with shaking. This culture was collected (10 mL, t0) and used to inoculate replicate 4 mL LB cultures (+/- 1mM IPTG and antibiotics) at 0.01 *A*_600_, which were then repeatedly grown 130 min to 0.3 *A*_600_ (5x doublings) and back-diluted to 0.01 for a total of 3 times (15x doublings). At the endpoint, cultures were collected (4 mL, t15) by pelleting (9000 × g for 2 min) and stored at −80° C. The following day, genomic DNA was extracted using the DNeasy Blood & Tissue kit (Qiagen #69506) with the recommended pre-treatment for Gram-negative bacteria and a RNAse A treatment. sgRNA spacer sequences were amplified from gDNA using Q5 polymerase (NEB) for 14x cycles using custom primers containing TruSeq adapters and indices, followed by gel-purification from 8% TBE gels. All sequencing was performed at the Chan Zuckerberg Biohub on the Illumina NextSeq platform using Single End 50bp reads.

Design of the sequencing libraries was optimized to enable multiplexing of many samples and to ensure diversity during cluster generation on the Illumina platform. Custom primers were used to generate the sequencing library that incorporated a second barcode (4bp) to be read in Read 1 (SeqLib.A.F and SeqLib.B.F in Fig. S1). In combination with the TruSeq barcode incorporated by the opposite primer (SeqLib.R in Fig. S1), this enabled the samples to be effectively dual-indexed. In preparing the sequencing libraries, samples were split into two types, Library Type A and Library Type B, which differ in the offset position of the TruSeq Read 1 primer used for sequencing (Fig. S1). Briefly, Library Type A introduces a 2bp offset, as such that, when libraries of Type A and Type B are sequenced on the same flowcell, diversity of sequencing reads is ensured throughout the read length, which includes the spacer region (variable sequence) and the promoter region (identical sequence).

Spacer sequences were extracted from FASTQ files and counted by exact matching to expected library spacers. For each treatment condition, the counts tables from each biological replicate were used as inputs for DESeq2 to calculate the change in abundance (Log2FC) and statistical significance (*P*_value_ and adjusted *P*_value_).

### Determination of LogP values

LogP values (partition coefficient) were determined using Chemicalize from ChemAxon (Escondido, California, USA).

### Preparation of graphs

GraphPad Prism software (v.7.00) was used to generate graphs and performed statistical tests. Number of independent experiments are indicated in each figure legend.

## RESULTS

### Susceptibility of bacteria to CAMPs and ceragenins

Minimal inhibitory concentrations (MIC) for the CAMPs colistin and LL37 as well as two ceragenin compounds, CSA13 and CSA131 (see structures in FIG. 1A), were determined against the Gram-negative bacterium *E. coli*, the Gram-positive bacterium *L. monocytogenes* and several mycobacterial species (i.e. *M. avium, M. marinum, M. smegmatis* and *M. tuberculosis*) (FIG. 1B-1H and Table S1). The fluoroquinolone antibiotic ciprofloxacin (CIP, which inhibits DNA gyrase) was included as a positive control. As expected, colistin, which requires binding to LPS for activity (35), was active against *E. coli* (FIG. 1B), but not against *L. monocytogenes* (FIG. 1C) and each of the mycobacterial species (FIG. 1D–1H). Interestingly, LL37 was active against *E. coli* (FIG. 1B) and *L. monocytogenes* (FIG. 1C) but had no detectable activity against mycobacterial species (FIG. 1D–1H). The ceragenins CSA13 and CSA131 were also active against both *E. coli* (FIG. 1B) and *L. monocytogenes* (FIG. 1C). In contrast to colistin and LL37, the ceragenins had activity against mycobacteria, although the MICs varied between species (FIG. 1D–1H). While *M. smegmatis* was highly susceptible to CSA13 and CSA131 (FIG. 1G), both compounds were less active against the slower-growing species *M. avium* (FIG. 1D and 1E), *M. marinum* (FIG. 1F) and *M. tuberculosis* (FIG. 1H). Similar trends in MIC values for *E. coli, L. monocytogenes* and *M. smegmatis* were observed with two other ceragenin compounds, CSA44 and CSA144 (See Table S1), which further confirmed that ceragenins have antimicrobial activity against mycobacteria. Overall, these results demonstrate that the spectrum of activity of ceragenins is broader than colistin and LL37, indicating different requirements for activity.

**FIGURE 1.**
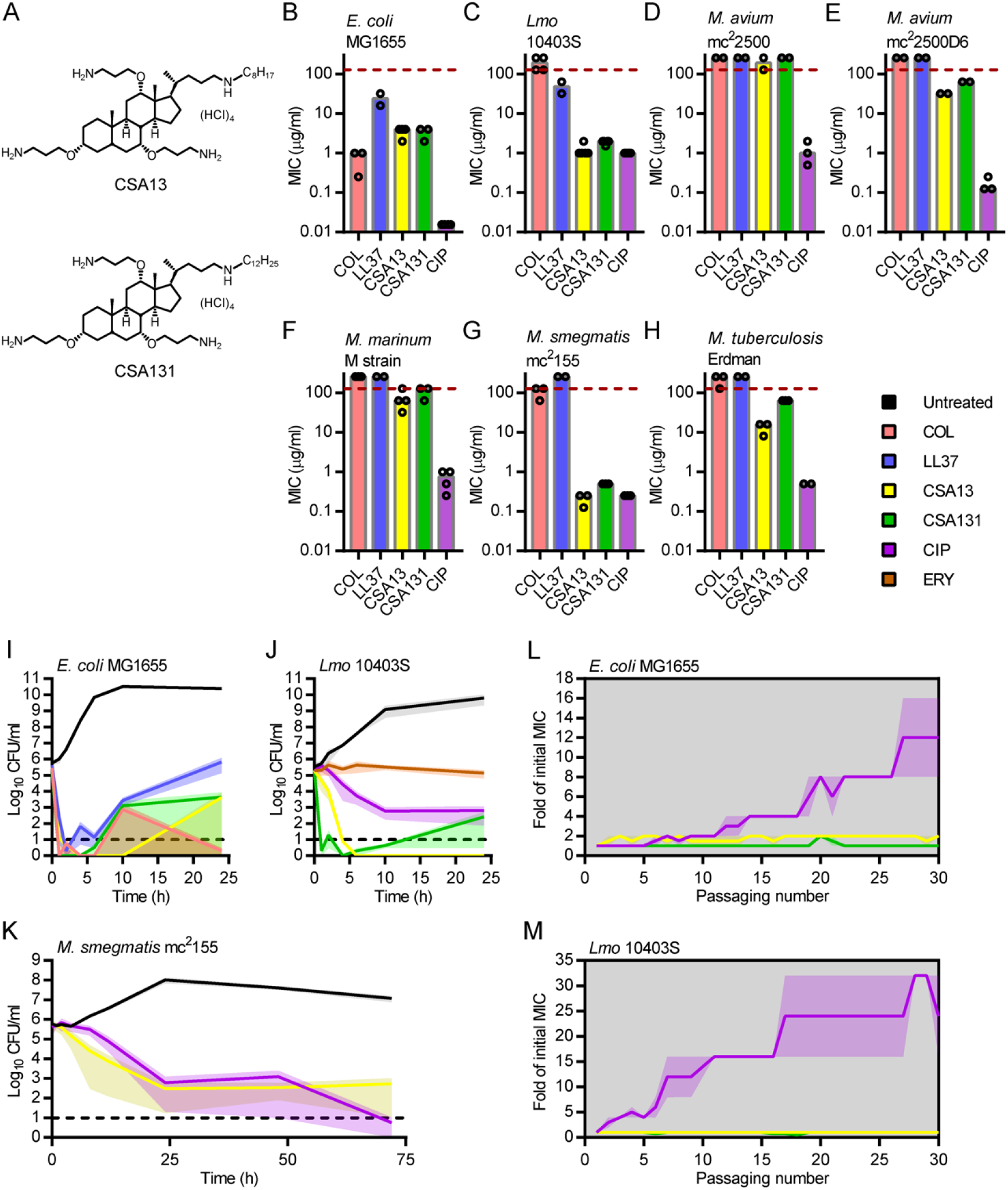
Ceragenins kills phylogenetically diverse bacteria. (A) Structures of the ceragenins CSA13 and CSA131. Minimal inhibitory concentrations (MIC) of colistin (COL), LL37, CSA13, CSA131 and ciprofloxacin (CIP) against *E. coli* MG1655 (B), *L. monocytogenes* (Lmo) 10403S (C), *M. avium* mc^2^2500 (D), *M. avium* mc^2^2500D6 (E), *M. marinum* M strain (F), *M. smegmatis* mc^2^ 155 (G) and *M. tuberculosis* Erdman (H). Dots and bars indicate results from independent experiments and median values, respectively. Time-kill experiments of *E. coli* MG1655 (I), *L. monocytogenes* 10403S (J) and *M. smegmatis* mc^2^ 155 (K) exposed to colistin (in red), LL37 (in blue), CSA13 (in yellow), CSA131 (in green), ciprofloxacin (in purple) and/or erythromycin (ERY; in orange), a bacteriostatic antibiotic. Untreated samples are in black and results are showed as means of two independent experiments. Shaded areas show standard error of the mean (SEM). Serial passages of *E. coli* (L) and *L. monocytogenes* (M) exposed to CSA13 (in yellow), CSA131 (in green) and ciprofloxacin (in purple). Bacteria were passaged daily in presence of sub-inhibitory concentrations of antibiotics. Results are expressed as means and SEM of two independent experiments.

### Ceragenins are bactericidal

To determine if ceragenins kill all three types of bacteria, we performed kill-curve experiments at inhibitory concentrations (~1-2 × MICs) of the molecules (FIG. 1I–1K). Ceragenins (CSA13 or CSA131) killed *E. coli* (FIG. 1I), *L. monocytogenes* (FIG. 1J) and *M. smegmatis* (FIG. 1K), although some bacterial cultures recovered during this time course. These results confirmed that ceragenins act on bacteria through a bactericidal mechanism.

### Serial passage of *E. coli* and *L. monocytogenes* in the presence of sub-inhibitory concentrations of ceragenins

Isolation and characterization of antibiotic-resistant bacteria could provide insight into the mode of action of ceragenins. As such, the generation of ceragenin-resistant bacteria was attempted by performing serial passaging experiments with *E. coli* and *L. monocytogenes* in the presence of sub-inhibitory concentrations of ciprofloxacin, CSA13 and CSA131 (FIG. 1L and 1M). In contrast to ciprofloxacin-exposed bacteria, *E. coli* and *L. monocytogenes* bacteria exposed to ceragenins did not give rise to stable resistance. The generation of spontaneous *M. smegmatis* mutants resistant to CSA13 was also attempted, but no CSA13-resistant bacteria were recovered, although bacteria resistant to ciprofloxacin and rifampicin were isolated from parallel control experiments (data not shown). These results confirmed that resistance to ceragenins is infrequent (14) and does not emerge *in vitro* under conditions known to generate resistant mutants against antibiotics.

### Transcriptional response of *E. coli* exposed to ceragenins

The transcriptional response of bacteria to antibiotics was analyzed to gain insights into the mechanism of action of CAMPs and ceragenins, as similarly reported for other antibacterial compounds (36, 37). More specifically, we determined the global transcriptional responses of *E. coli* exposed to colistin, LL37, CSA13 and CSA131 using RNAseq. Bacteria were grown to log phase and treated with supra-MIC concentrations of antibiotics for one hour before being harvested for RNA extraction and sequencing (see Materials and Methods). Plots of the normalized number of reads per gene showed an excellent correlation (R = 0.965-0.999) between biological replicates for each of the conditions tested (FIG. 2A–2D), demonstrating the reproducibility of the method. Hundreds of statistically significant changes in gene expression (defined by absolute log_2_ fold change >1 and adjusted *P*_value_ < 0.05) following exposure of bacteria to colistin, LL37, CSA13 and CSA131 were measured (FIG. 2E–2H and Data Set S1). These results validated our RNAseq approach for the analysis of the transcriptional response of *E. coli* to antibiotics.

**FIGURE 2.**
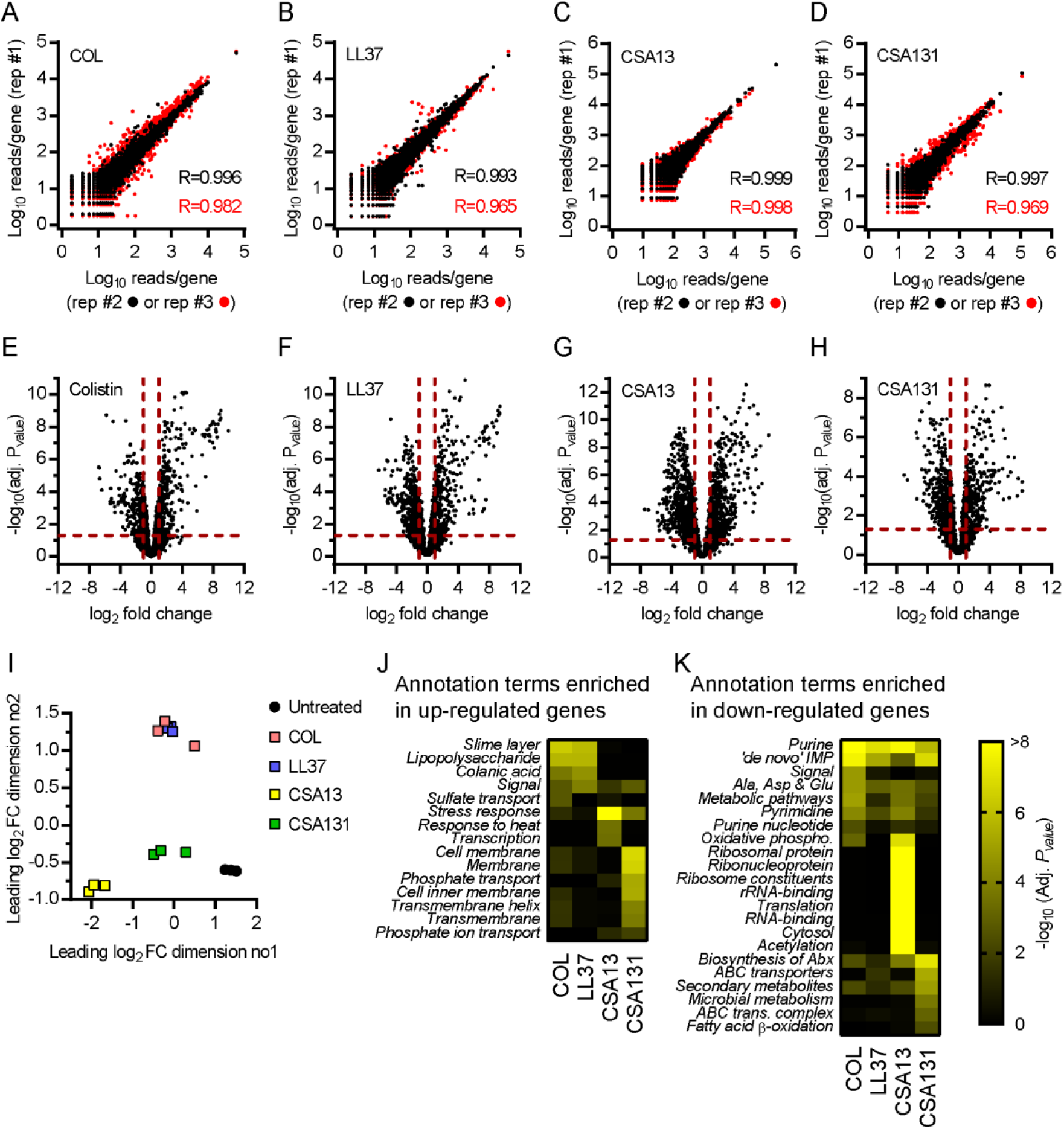
Transcriptomic response of *E. coli* exposed to ceragenins. RNA from exponentially growing *E. coli* bacteria exposed to supra-MIC concentrations of colistin (COL), LL37, CSA13 and CSA131 was extracted and sequenced. (A-D) Replica plots showing the log_10_ of normalized number of reads per gene for bacteria exposed to antibiotics. Correlation coefficients (R) between replicates #1 and #2 (●) and #1 and #3 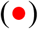 are displayed. (E-H) Volcano plots that represent RNA expression as means of log_2_ fold changes and −log_10_ adjusted *P*_values_ (adj. *P*_value_) for bacteria exposed to antibiotics in comparison to untreated control samples. Horizontal and vertical dotted red lines indicate adjusted *P* values less than 0.05 (or −log_10_ (adj. *P*_value_) greater than 1.3) and absolute log_2_ fold changes greater than 1. (I) Multidimensional scaling (MDS) plot showing the separation between biological replicates and between untreated and antibiotic-treated samples. (J) Annotation terms enriched for genes significantly up-regulated (log_2_ FC > 1 and adj. *P*_value_ < 0.05) following exposure to antibiotics. (K) Annotation terms enriched for genes significantly down-regulated (log_2_ FC < −1 and adj. *P*_value_ < 0.05) following exposure to antibiotics. Adjusted *P*_values_ of annotation terms associated with a false discovery rate (FDR) value > 0.05 for at least one antibiotic are showed. Only the 8 most statistically significant annotation terms are showed for each conditions. Annotation terms are abbreviated and/or modified for a purpose of presentation (see Data Set S2 for a more detailed information). Data are from 3 independent experiments.

The global transcriptional responses of *E. coli* in response to CAMPs and ceragenins was analyzed using a multidimensional scaling analysis (FIG. 2I). Interestingly, while transcriptional responses of bacteria to CAMPs were extremely similar, the response to CSA13 and CSA131 were not only distinct from these CAMPs, but also distinct from each other. Those trends were corroborated using pathway analysis that showed the enrichment of annotation terms associated with the outer membrane (e.g. lipopolysaccharide and colanic acid) in genes up-regulated by CAMPs, but not ceragenins, (FIG. 2J and Data Set S2) as well as the enrichment of terms associated with translation in genes down-regulated by CSA13, but not CSA131 (FIG. 2K and Data Set S2). While the two ceragenins are structurally quite similar, the addition of four methylene groups to the CSA13 carbon chain to create CSA131 significantly increases the hydrophobicity of the molecule, increasing the partition coefficient from LogP_CSA13_ = 5.51 to LogP_CSA131_ = 7.29 (see Materials and Methods), which likely contributes to differences in antibacterial activities. Overall, these results showed that transcriptional responses of *E. coli* to the naturally-occurring CAMPs colistin and LL37 are similar, but differ from the response to ceragenins. These results also suggested that *E. coli* responds differently to the structurally related ceragenin compounds, CSA13 and CSA131.

### Identification of pathways defining the transcriptional response of *E. coli* to ceragenins

Pathway analysis of genes modulated by more than one compound was performed to further define transcriptional responses to CAMPs and ceragenins. The Venn diagram in Figure 3A visualizes the extent of overlap of the individual *E. coli* genes that had significant increases in mRNA abundance upon treatment with each of the molecules. In particular, 86 genes were induced in all four conditions, 68 were upregulated specifically during CAMP treatment while 57 were induced by the ceragenins (FIG. 3A and Data Set S3). The annotation term “signal” was significantly enriched among the genes up-regulated by all antibiotics (FIG. 3A and Data Set S3), which might be indicative of a common response to CAMPs and ceragenins.

**FIGURE 3.**
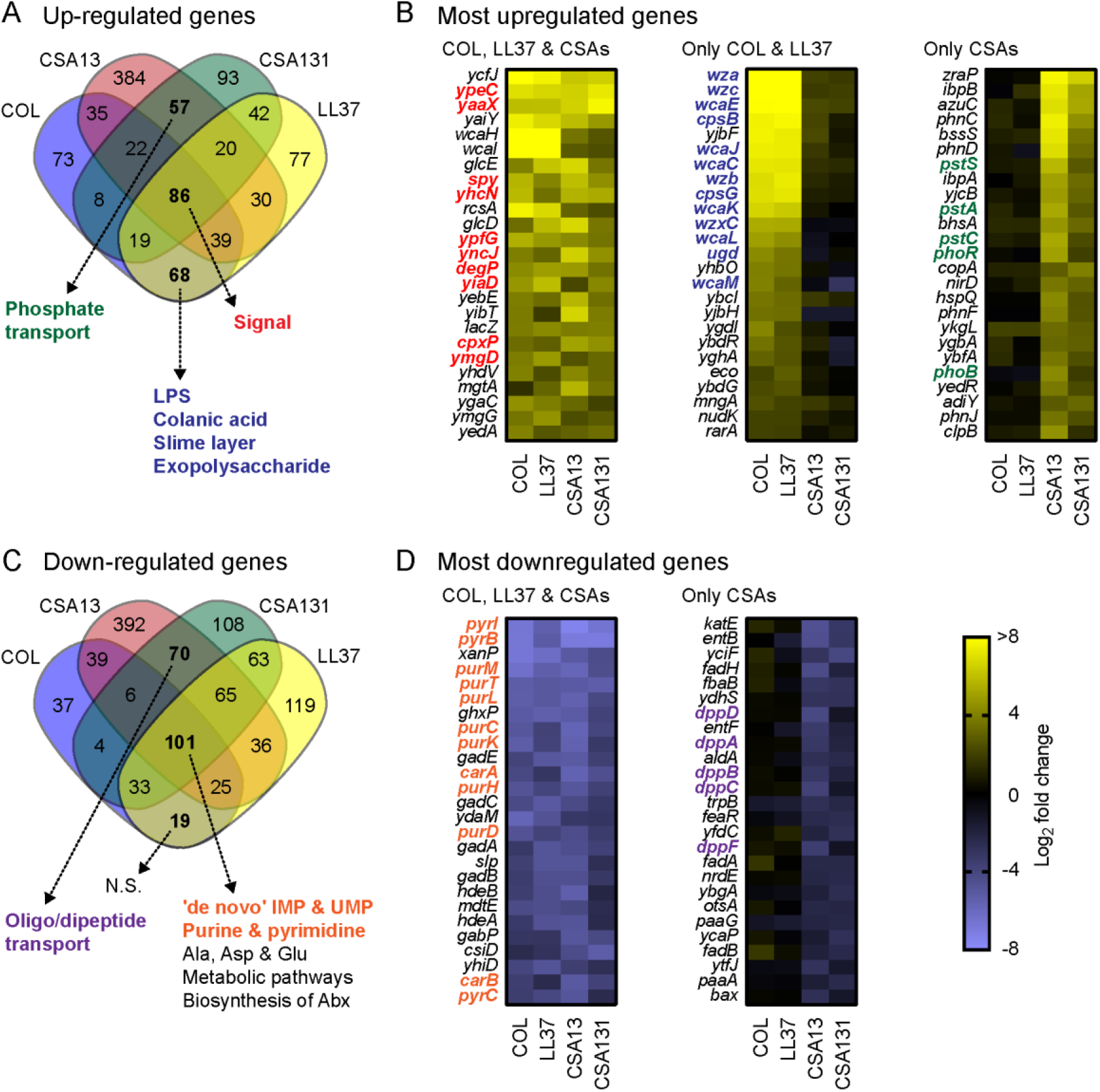
Identification of pathways defining the transcriptional response of *E. coli* to ceragenins. (A) Venn diagram analysis of genes significantly up-regulated (log_2_ FC > 1 and adjusted *P*_value_ < 0.05) in *E. coli* exposed to antibiotics. Annotation terms associated with a false discovery rate (FDR) value > 0.05 for genes up-regulated by all antibiotics, by CAMPs or by ceragenins are shown by dotted arrows. (B) Top 25 most up-regulated genes for bacteria exposed to all antibiotics, to CAMPs or to ceragenins. Genes belonging to the annotation terms «signal» (in red), «LPS», «colanic acid», «slime layer» and «exopolysaccharide» (in blue) and «phosphate transport» (in green) are indicated. (C) Venn diagram analysis of genes significantly down-regulated (log_2_ FC < −1 and adjusted *P*_value_ < 0.05) in *E. coli* exposed to antibiotics. Annotation terms associated with a false discovery rate (FDR) value > 0.05 for genes down-regulated by all antibiotics, by CAMPs or by ceragenins are shown by dotted arrows. (D) Top 25 most down-regulated genes for bacteria exposed to all antibiotics or to ceragenins. Genes belonging to the annotation terms «de novo IMP», «de novo UMP», «purine» and «pyrimidine» (in orange) and «oligo/dipeptide transport» (in purple) are indicated. Annotation terms are abbreviated and/or modified for the purpose of presentation (see Data Set S3 for a more detailed information). Data are from 3 independent experiments.

Interestingly, this group included genes involved in the membrane stress response such as *spy*, *degP* and *cpxP* (38) (Fig. 3B). Consistent with results from FIG. 2, genes specifically up-regulated in bacteria exposed to CAMPs were significantly associated with annotation terms related to LPS/colanic acid biosynthesis and included genes such as *wzc, wcaE* and *cpsB* (FIG. 3A, 3B and Data Set S3). Interestingly, genes specifically up-regulated in bacteria exposed to ceragenins were significantly associated with the annotation term “phosphate transport” (FIG. 2J, FIG. 3A and Data Set S3) and include genes encoding the major phosphate-responsive regulators PhoR and PhoB (FIG. 3B). Overall, these results suggested that *E. coli* responds to CAMPs and ceragenins by upregulating genes involved in signaling and response to membrane stress. These results also showed that while exposure of *E. coli* to CAMPs induces the expression of genes related to LPS/colanic acid biosynthesis, exposure to ceragenins induces the expression of genes involved in phosphate transport.

For genes with mRNA levels that decreased during these treatments, 101 were down-regulated by all antibiotics, 19 by CAMPs, and 70 by ceragenins (FIG. 3C and Data Set S3). Pathway analysis identified enrichment of terms related to amino acids and nucleotide metabolism in genes down-regulated following exposure to all antibiotics (FIG. 3C and Data Set S3). This finding was corroborated by the finding that genes related to purine and pyrimidine biosynthesis (e.g. *pyrB, purM* and *purT*) were among the most significantly down-regulated genes by these molecules (FIG. 3D), which is consistent with our pathway analysis (FIG. 2K). Whereas no annotation terms were significantly enriched for genes only down-regulated by CAMPs, genes down-regulated by ceragenins showed an enrichment for genes involved in oligopeptide/dipeptide transport such as *dppD* and *dppA* (FIG. 3C, FIG 3D and Data Set S3). Although the reason for the downregulation of genes involved in peptide transport in *E. coli* exposed to ceragenins is unknown, these results suggested that bacteria exposed to CAMPs and ceragenins respond by downregulating genes involved in metabolic pathways.

### Identification of cis-acting elements that regulate the transcriptional response of *E. coli* to ceragenins

To determine which signal transduction pathways control the transcriptional responses to CAMPs and ceragenins, the DNA sequences immediately 5’ of genes with significantly altered mRNA levels were analyzed for the presence of cis-acting promoter and operator sequences known or predicted to recruit transcription factors (29). Interestingly, genes up-regulated following exposure to each of the molecules (FIG. 4A and Data Set S4) were associated with cis-acting elements interacting with the response regulator CpxR of the CpxA/CpxR two-component regulatory system (enrichment of 11.63%; 10 out of 86 genes), which responds to envelope stress (39) and is consistent with the upregulation of the CpxR-regulon genes *spy*, *degP* and *cpxP* (FIG. 3B). This analysis also showed enrichment for genes associated with cis-acting elements binding the primary sigma factor σ^D^ (40), also involved in the redistribution of the RNA polymerase in response to osmotic stress (41), the alternative sigma factor σ^E^ that coordinates the envelope stress response (39, 42, 43), and the alternative sigma factor σ^H^, which controls the expression of heat shock genes as well as genes involved in membrane functionality and homeostasis (44) (FIG. 4A). Genes down-regulated following exposure to all antibiotics were associated with the presence of binding sites for the HTH-type transcriptional repressor PurR (FIG. 4B and Data Set S4), which regulates genes involved in the de novo synthesis of purine and pyrimidine nucleotides (45, 46) and corroborates our above analysis (FIG. 2K, 3C and 3D). Overall, these results suggested that CAMPs and ceragenins perturb the bacterial envelope and trigger the CpxA/CpxR system. These results also suggested that PurR down-regulates the expression of genes involved in the biosynthesis of purine and pyrimidine following the exposure of *E. coli* to CAMPs and ceragenins.

**FIGURE 4.**
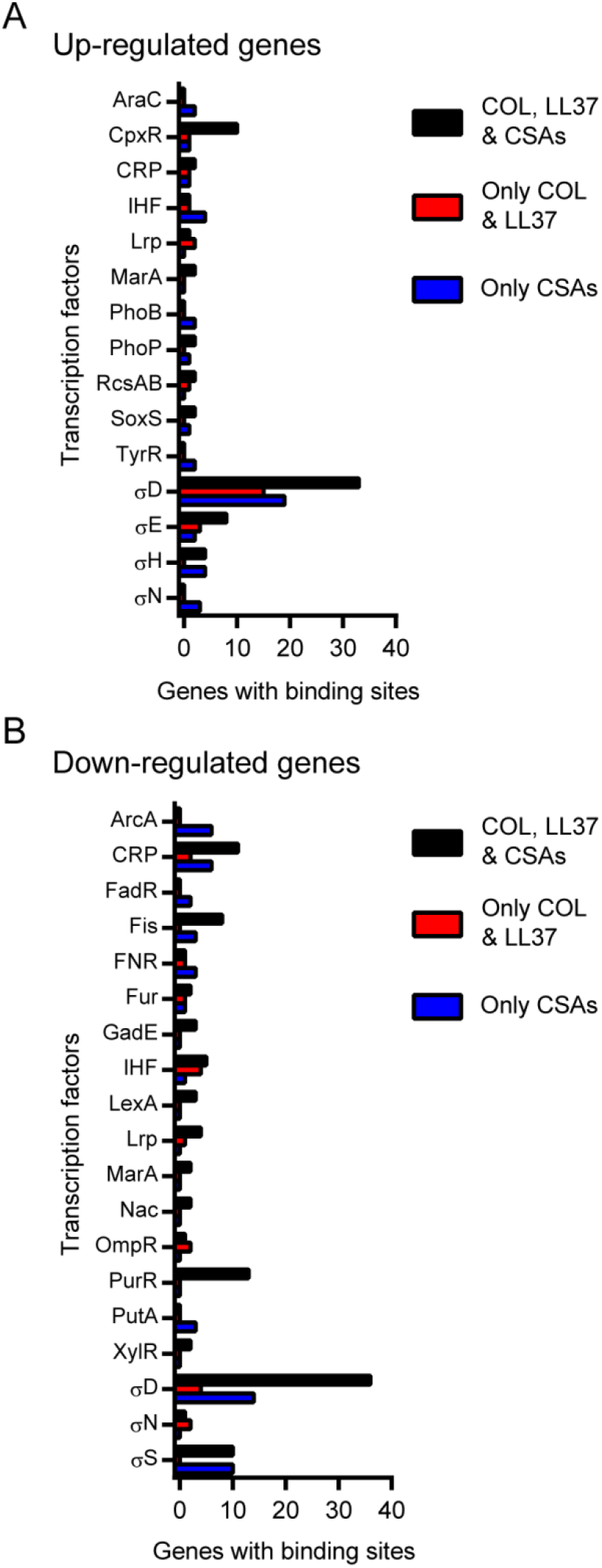
Identification of cis-acting elements that regulate the transcriptional response of *E. coli* to ceragenins. Genes with known or predicted promoters or binding sites for transcription factors are enumerated for genes commonly up-regulated (A) or down-regulated (B) by all antibiotics (black bars), by CAMPs (red bars) or by ceragenins (in blue). Only promoters/binding sites identified more than once for at least one group are represented.

### Expression of the CpxR, PurR, RcsA and PhoB regulons in *E. coli* exposed to ceragenins

To further confirm a role for CpxR and PurR in the response of *E. coli* to CAMPs and ceragenins, the expression of the CpxR and PurR regulons was analyzed in more detail (FIG. 5A-B and Data Set S4). The heat map of the CpxR regulon showed a consistent regulation of several genes in bacteria exposed to all four compounds (e.g. *cpxP, degP, dsbA* and *spy*) (FIG. 5A) and corroborates our results described above (FIG. 3B and 4A). Also consistent with our findings described above (FIG. 2K, 3C, 3D and 4B), the heat map of the PurR regulon (FIG. 5B) showed downregulation of most genes reported to be repressed by this transcription factor. These results confirmed that the expression of the CpxR and PurR regulons are modulated in *E. coli* exposed to CAMPs and ceragenins.

**FIGURE 5.**
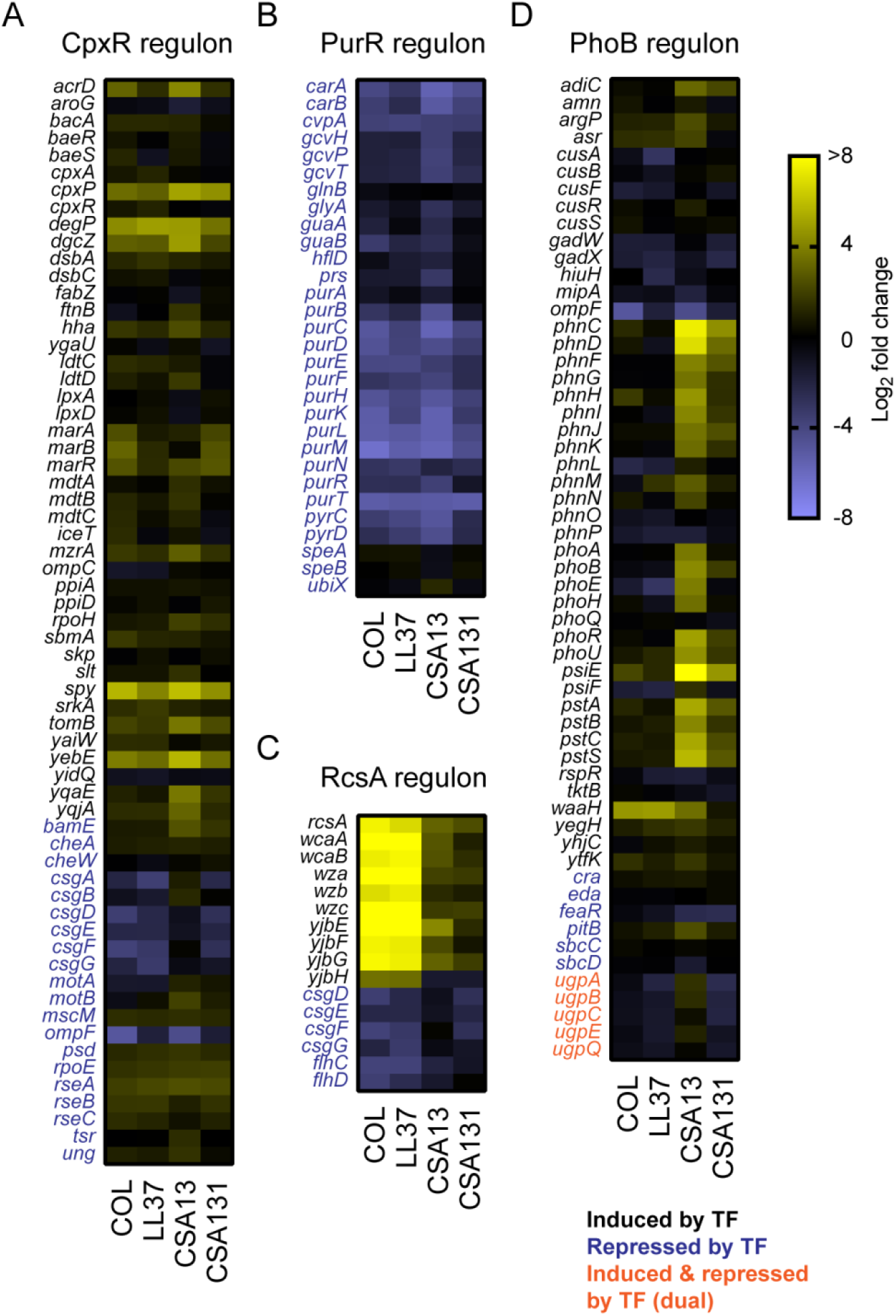
Expression of the CpxR, PurR, RcsA and PhoB regulons in *E. coli* exposed to ceragenins. Heat maps of log_2_ fold changes for genes of the CpxR (A), PurR (B), RcsA (C) and PhoB (D) regulons in bacteria exposed to antibiotics. Genes predicted to be induced (in black), repressed (in blue) or both (in orange) by a specific transcription factors are indicated. Data are from 3 independent experiments.

Our expression analysis led us to also focus on the RcsA and PhoB regulons to gain insight into the differential regulation of genes involved in the biosynthesis of colonic acid and phosphate transport, respectively (FIG. 5C-D and Data Set S4). RcsA regulates the expression of genes involved in colanic acid biosynthesis (39), a pathway that was up-regulated in *E. coli* exposed to CAMPs, but not ceragenins (FIG. 2J, 3A and 3B). Accordingly, the heat map of the RcsA regulon showed a marked modulation of this pathway in *E. coli* exposed to CAMPs in comparison to ceragenins (FIG. 5C). The phosphate regulon transcriptional regulatory protein PhoB regulates the expression of genes involved in phosphate transport (47) and is up-regulated in *E. coli* following exposure to ceragenins, but not CAMPs (FIG. 2J, 3A and 3B). Accordingly, the heat map of the PhoB regulon showed partial but specific induction in bacteria exposed to ceragenins (FIG. 5D). More specifically, upregulation of the *phn C-phnP* operon, the *pstSCAB-phoU* operon as well as a trend for other *pho* genes (e.g. *phoB* and *phoR*) and the phosphate starvation-inducible *psiEF* genes were specifically observed in bacteria exposed to ceragenins (FIG. 5D). These results suggested that the specific upregulation of genes involved in colanic acid biosynthesis and phosphate transport in bacteria exposed to colistin/LL37 and ceragenins are mediated by RcsA and PhoB, respectively.

### Proteomic response of *E. coli* exposed to colistin and CSA13

To determine if the changes in mRNA levels in response to the molecules led to changes in the proteome, we measured global protein abundance in bacteria exposed to colistin and CSA13 by mass spectrometry-based proteomics. *E. coli* cultures were grown to log phase and treated with supra-MIC concentrations of antibiotics before protein extraction, peptide preparation and peptide quantification (see Materials and Methods). Approximately 1800 unique proteins were detected for each biological replicate (FIG. 6A) and the number of unique peptides identified showed an excellent correlation between biological replicates (FIG. 6B). Several statistically significant changes in protein expression (absolute log_2_ fold change >1 and adjusted *P*_value_ < 0.05) were observed following exposure of *E. coli* to colistin (FIG. 6C) and CSA13 (FIG. 6D) (see Data Set S5). The dataset showed that *E. coli* exposed to both colistin and CSA13 up-regulated the proteins DegP, Spy and YepE, which are known members of the Cpx regulon (38) that were strongly up-regulated at the transcriptional level following exposure to CAMPs and ceragenins (FIG. 5A). The dataset also showed the modulation of proteins involved in colanic acid biosynthesis (e.g. Ugd and WcaG) or related to the PhoB regulon (e.g. PstB and PstS) in bacteria exposed to colistin or CSA13, respectively, which also corroborate our transcriptional data (FIG. 5C and 5D). Then, similarly to our transcriptional analysis, the proteomic data showed a common induction of the Cpx envelope stress response, but also the specific induction of proteins involved in colanic acid biosynthesis by colistin as well as the modulation of members of the PhoB regulon by CSA13.

**FIGURE 6.**
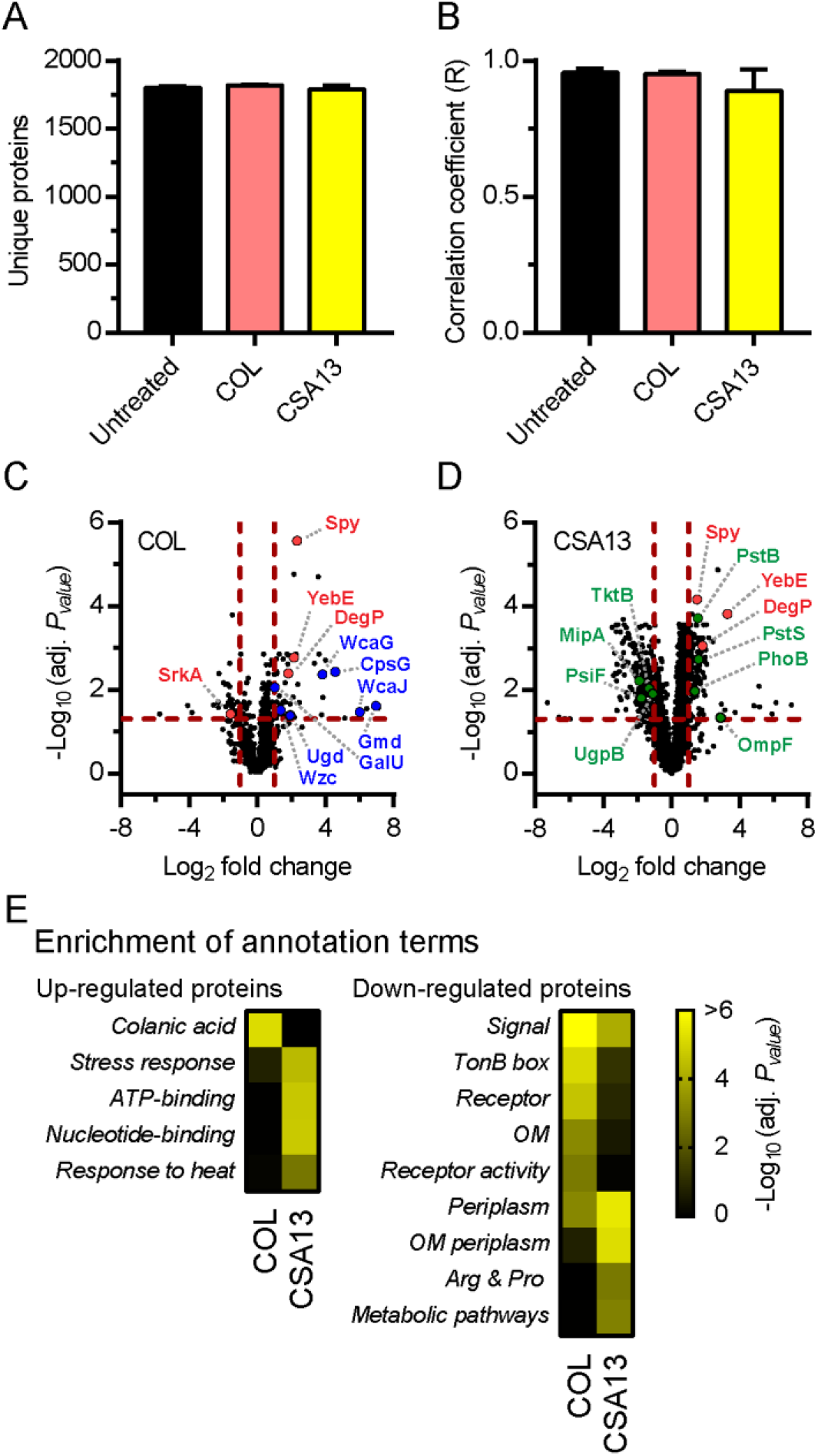
Proteomic response of *E. coli* exposed to colistin and CSA13. (A) Number of unique proteins identified for each conditions. (B) Correlation coefficient (R) between the number of peptides per proteins between the biological replicates of untreated bacteria and bacteria exposed to colistin (COL) or CSA13. Data are represented as means and standard deviations. (C-D) Volcano plots that represent protein expression as means of log_2_ fold changes and −log_10_ adjusted *P*_values_ (adj. *P*_value_) for bacteria exposed to antibiotics in comparison to untreated controls. Horizontal and vertical dotted red lines indicate adjusted *P* values less than 0.05 (or −log_10_ (adj. *P*_value_) greater than 1.3) and absolute log_2_ fold changes greater than 1. Some proteins that are members of the Cpx regulon (in red), involved in colanic acid biosynthesis (in blue) or members of the PhoB regulon (in green) are highlighted. (E) Annotation terms enriched for proteins significantly up- or down-regulated (absolute log_2_ FC > 1 and adj. *P*_value_ < 0.05) following the exposure of *E. coli* to colistin and CSA13. Adjusted *P*_values_ of annotation terms associated with a false discovery rate (FDR) value > 0.05 for at least one antibiotic are showed. Annotation terms are abbreviated and/or modified for the purpose of presentation (see Data Set S6 for a more detailed information). Data are from 3 independent bacterial cultures for each conditions.

Annotation term enrichment analysis was performed on the proteomic dataset in order to identify pathways significantly modulated in bacteria exposed to colistin and CSA13 (see Data Set S6). Similar to our transcriptional results (FIG. 2J), the colanic acid pathway was significantly enriched among proteins up-regulated by colistin, but not CSA13 (FIG. 6E). A heat response signature was significantly enriched among proteins uregulated by CSA13, but not colistin (FIG. 6E), corroborating the transcriptional upregulation of genes associated with cis-acting elements for σ^H^ (FIG. 4A). In addition, while pathways associated with the periplasm were enriched among proteins down-regulated by both colistin and CSA13, the outer membrane pathway was significantly enriched among proteins down-regulated by colistin, but not by CSA13 (FIG. 6E). Overall, these results confirmed that *E. coli* responds distinctly to colistin and CSA13, although both compounds modulated proteins associated with the bacterial envelope.

These results also confirmed findings from the transcriptional analysis and showed that colistin and CSA13 modulate the colanic acid and the response to heat pathways, respectively.

### Identification of genetic determinants of resistance to ceragenins in *E. coli*

Our inability to identify ceragenin-resistant *E. coli* mutants (FIG. 1L and 1M) is consistent with results previously published by Pollard *et al*. (14), indicating that resistance to ceragenins emerges infrequently in culture despite strong selective pressure. This suggests that ceragenins either have multiple essential targets or affect cellular structures that are immutable. Thus, traditional genetic approaches to identify the target(s) of ceragenins has not been feasible. In order to gain insight into the genetic determinants of ceragenin action, we employed an alternative genetic approach that utilizes Clustered Regularly Interspaced Short Palindromic Repeats interference (CRISPRi) to reduce expression of genes in *E. coli* (FIG. 7A). As demonstrated previously, this approach allows for partial knockdown of essential *E. coli* genes, practically creating hypomorphic alleles that have reduced function but still promote cell viability (48, 49). By combining these genetic perturbations with subinhibitory concentrations of antibiotics and by measuring the effects on bacterial fitness, we sought to identify functional interactions between bacterial pathways and antibiotic stress.

**FIGURE 7.**
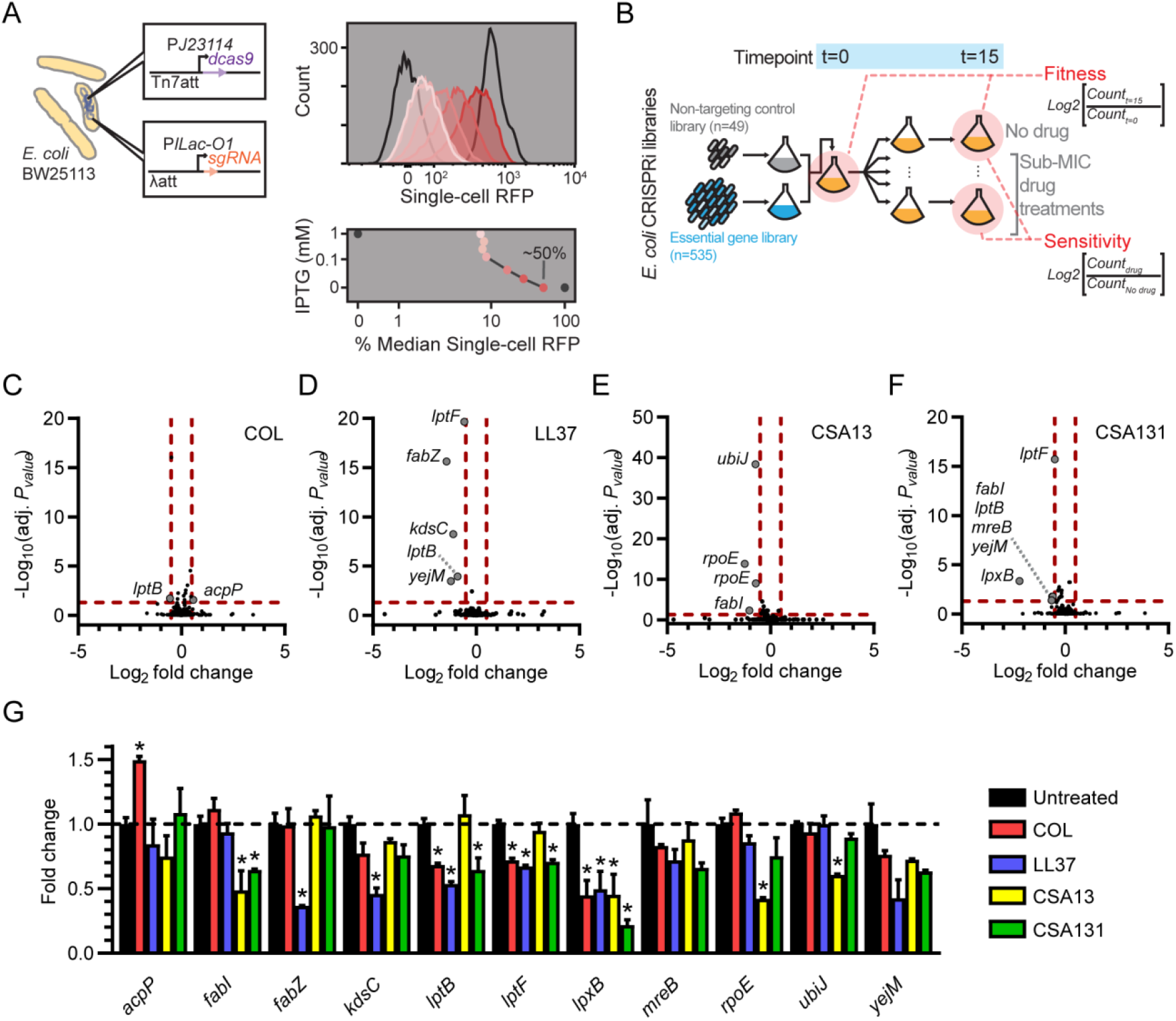
Identification of the genetic determinants of resistance to ceragenins in *E. coli*. (A) Details and calibration of the *E. coli* CRISPRi system. dCas9 and sgRNA expression cassettes were integrated into the chromosome (Tn7att and lambda att, respectively) and controlled by weak constitutive (dcas9) or inducible (sgRNA) promoters, as indicated. Right panels show that CRISPRi produces unimodal reduction in expression when targeting chromosomal *rfp* (i.e. a gene encoding a red fluorescent protein). Median percentages of knockdown are indicated in the right lower panel. (B) Schematic of the pooled growth experiments of *E. coli* CRISPRi libraries in the presence or absence of antibacterial compounds. The strain-specific metrics of fitness and sensitivity (drug-specific) are calculated as the change in relative abundance of each strain between the two time points or conditions, using the formulae as indicated. Changes in abundance (Log_2_FC) and adjusted *P_values_* (-Log_10_(adj. *P_value_*)) associated with each strain following exposure to colistin (COL) (C), LL37 (D), CSA13 (E) and CSA131 (F) are shown. Horizontal and vertical dotted red lines indicate adjusted *P* values less than 0.05 (or −log_10_ (adj. *P*_value_) greater than 1.3) and absolute log_2_ fold changes greater than 0.5. Genes associated with significant changes in abundance are labeled for each compounds. (G) Mean fold changes in abundance and standard deviations (SD) associated with significantly enriched or depleted CRISPRi strains (only one *rpoE-* targeting strain is shown) following exposure to COL, LL37, CSA13 and CSA131 (*, *P* < 0.05 [Two-tailed unpaired *t*-test]). Means and SDs were calculated from counts normalized to the total number of counts for each conditions. Data are from two biological replicates.

We screened a pooled library of inducible essential gene knockdown strains grown with or without subinhibitory concentrations of CAMPs or ceragenins (FIG. 7B). To evaluate the fitness of each individual CRISPRi knockdown within the complex population, we used deep sequencing to measure the relative abundance of guide sequences during the different culturing conditions (see Materials and Methods and FIG. S1). Fold changes in abundance (Log_2_FC) in response to CAMPs and ceragenins were calculated in comparison to the control condition for each knockdown strain, and significantly resistant or sensitized strains (defined by absolute log_2_ fold change > 0.5 and an adjusted *P*_value_ < 0.05) were identified for each of the compounds (FIG. 7C–7F and Data Set S7). Interestingly, the CRISPRi guides in the sensitized or enriched strains for one or several treatments predominantly targeted genes involved in the bacterial envelope, consistent with our earlier results that both CAMPs and ceragenins affect the bacterial envelope.

The changes in abundance for the genes identified above were further analyzed and compared between treatments (FIG. 7G). Genes involved in the LPS biosynthetic pathways were predominant in these screens, and knockdown of lipid-A-disaccharide synthase LpxB (50) sensitized *E. coli* to all the compounds, highlighting again the bacterial surface as a common site of action for CAMPs and ceragenins. Silencing of the LPS transport genes *lptB* and *lptF* (51) led to sensitivity to colistin, LL37 and CSA131, but not to CSA13, which corroborates the above transcriptomic data (FIG. 2I-K) and again suggests distinctive mechanisms of action for CSA13 and CSA131. Furthermore, knockdown of *kdsC* (56), encoding an enzyme involved in LPS biosynthesis, also led to sensitivity to LL37. Knockdown strains for *rpoE*, the gene encoding the envelope stress responsive sigma factor σ^E^ (39, 42, 43), and the ubiquinone biosynthesis gene *ubiJ* (58) were sensitive to CSA13, but not to CSA131, also supporting the idea that these ceragenins have distinctive mechanisms of action. On the other hand, the knockdown strain for the fatty acid biosynthesis gene *fabI* (52, 53) was sensitive to both CSA13 and CSA131 and not to the CAMPs, and may constitute a common molecular determinant of sensitivity to ceragenins. Other genes involved in fatty acid metabolism were identified as genetic interactors with CAMPs. More specifically, interference with the expression of *acpP*, encoding acyl carrier protein (54), led to resistance to colistin, whereas knockdown of *fabZ*, encoding a lipid dehydratase (55), led to sensitivity to LL37, demonstrating differences between the mechanisms of action of these two CAMPs, as previously suggested (35, 57). Taken together, these results support the notion that both CAMPs and ceragenins work by similar yet distinct mechanisms. These studies also provide a starting point for genetic determination of the mode of action for ceragenins.

## DISCUSSION

Although ceragenins were originally designed to mimic the physiochemical properties of CAMPs (11, 12), our results indicate that they evoke different responses from bacteria than naturally-occurring CAMPs. The fact that they work on a broader array of microbes than CAMPs, including mycobacteria, as showed by this study, also suggests that they have different mechanisms of action. Our results showed that ceragenins target an essential and conserved feature of the cellular envelope and kill phylogenetically diverse bacteria. Using trancriptomics, proteomics and a CRISPRi genetic approach, we compared the responses of bacteria to CAMPs and ceragenins and revealed similarities, but also striking differences, and showed that ceragenins trigger a distinctive envelope stress response. Interestingly, our data also suggested that the two prototypical ceragenins, CSA13 and CSA131, trigger different responses in bacteria. Overall, while our results confirmed that ceragenins act on the bacterial envelope, they challenged the assumption that CAMPs and ceragenins share the same mechanism of action.

Although ceragenins have the ability to kill mycobacteria, their activity varies considerably among species. The physicochemical properties of the mycobacterial cell envelope influences its permeability (59) and might explain the observed differences in susceptibility to ceragenins among mycobacterial strains and species (FIG. 1). The identification of the target(s) of ceragenins may reveal an essential feature of the clinically-relevant mycobacteria.

The profiling of the response of *E. coli* to CAMPs and ceragenins showed that these compounds trigger the Cpx envelope stress response, which is known to contribute to the bacterial adaptation to defects in the secretion and folding of inner membrane and periplasmic proteins (39, 60). This corroborates previous studies demonstrating that CpxR/CpxA influence the susceptibility of bacteria to CAMPs (61, 62) and suggests that the Cpx response might similarly help bacteria to survive exposure to ceragenins. The hypothesis that the envelope stress response is induced by CAMPs and ceragenins is also supported by the modulation of genes associated with cis-acting elements for σ^E^ (FIG. 4) and by the enrichment and/or depletion of CRISPRi strains targeting components of the bacterial envelope (FIG. 7).

A striking similarity between the transcriptomic profiles of bacteria exposed to CAMPs and ceragenins is the downregulation of genes involved in the biosynthesis of purines and pyrimidines (FIG. 2K, 3C, 3D, 4B & 5B). The cause of this downregulation is unknown, but it is possible that the repression of these metabolic pathways is part of the adaptive response to antibiotic exposure (63) and/or relates to a decrease requirement for nucleic acid in growth-inhibited bacteria. An intriguing question is whether the flux of the metabolites through these nucleotide metabolic pathways affect susceptibility to antimicrobial agents targeting the bacterial envelope, as observed for other antibiotics (64).

The results of this study showed that *E. coli* responds differently to CAMPs and ceragenins. We showed that CAMPs specifically induced the Rcs response and the expression of genes involved in the biosynthesis of colanic acid (FIG. 2J, 3A, 3B, 5C, 6C and 6E). This is consistent with a previous study that demonstrated that CAMPs, including polymyxin B and LL37, induce the Rcs regulon through the outer membrane lipoprotein RcsF (65). Surprisingly, the ceragenins CSA13 and CSA131 did not induce the Rcs response as markedly as CAMPs (FIG. 5C). In contrast with the current model that outer membrane perturbation by CAMPs is required for the activation of the Rcs response by RcsF (65, 66), our results show that ceragenins perturb the bacterial envelope of *E. coli* without extensively triggering the Rcs response. We also found that ceragenins, but not CAMPs, induced the expression of genes involved in phosphate transport and of the PhoB regulon (FIG. 2J, 3A, 3B, 5D and 6D). Although the reasons why these genes are differently modulated following exposure to antimicrobial compounds is not understood, these results strongly suggest that ceragenins and CAMPs might have distinctive mechanisms of action.

Despite some similarities between the response of bacteria exposed to ceragenins, such as the upregulation of several genes of the Cpx and PhoB regulons and the downregulation of genes involved in nucleotides metabolism, our data also showed striking differences between bacteria exposed to CSA13 and CSA131 (FIG. 2I, 2J, 2K and 7G). These differences include the upregulation of genes encompassing several functions (e.g. transcription factors and proteins involved in the heat response) as well as the downregulation of genes involved in protein translation in bacteria exposed to CSA13 (FIG. 2J and 2K). The cause of these differences is unknown, however, as noted above, the LogP values of CSA13 and CSA131 differ by almost 2 orders of magnitude. Given that the site of action is the bacterial envelope, such significant difference in coefficient partition values is likely to alter responses to membrane targets. Future work exploring the response of bacteria to a broader range of ceragenins will help in understanding these differences and might help in the design of compounds with a more defined mode of action.

Our CRISPRi approach identified sensitizing interactions between genes involved in the biology of the bacterial envelope and the antibacterial compounds colistin, LL37, CSA13 and CSA131 (FIG. 7). This information might prove valuable for the design of combination therapies that are synergistic and prevent the emergence of resistance, but also allow treatment regimens with lower concentrations of antibiotics and dose-related antibiotic toxicity (67–69). As an example, trilosan, a compound inhibiting the ceragenin-sensitivity determinant FabI (70) (FIG. 7G), might synergize with ceragenins. In addition, the CAMPs/ceragenins-sensitivity determinant LpxB was suggested as a target for the development of antibacterial compounds (71), which compounds would have the potential to more broadly synergize with CAMPs and ceragenins (FIG. 7G). Although those antibacterial interactions are purely speculative, our results suggest that the CRISPRi approach presented here constitutes a platform for target identification and the development of antibiotic combination therapies.

The results of this study suggested that CAMPs and ceragenins both kill bacteria by targeting the bacterial envelope. However, this study also supports the hypothesis that ceragenins have a distinctive mode of action and we propose a model in which ceragenins cross the outer layers of the bacterial envelope and disrupt the inner membrane. This hypothesis is supported by the broad spectrum of action of these molecules, which extend beyond bacteria. Whether the broader activity range of ceragenins impacts the selectivity for microbial membranes characteristic of endogenous CAMPs remains a key question for future study. A better understanding of the structure-activity relationship of these compounds and a deeper knowledge of their unique mechanism of action will be essential in the discovery of the next-generation of ceragenins with increased potency and selectivity.

## Supporting information

Data Set S1

Data Set S2

Data Set S3

Data Set S4

Data Set S5

Data Set S6

Data Set S7

## ACKNOWLEDGEMENTS

The authors would like to thank Teresa Repasy and Guillaume Golovkine for their assistance during the design of the RNAseq experiments, and Daniel A. Portnoy for providing *L. monocytogenes* 10403S. We acknowledge help from the UC Berkeley Functional Genomics Laboratory, the UC Davis Genome Center and the UC Davis Bioinformatics Core in performing and analyzing RNAseq experiments as well as the Chan Zuckerberg Biohub for the sequencing of the CRISPRi libraries. The UC Berkeley proteomics core provided assistance with the LC-MS analysis of our peptide samples. J.M.B was supported by NIH training grants (4T32HL007185-39 & −40; 5K12HL119997-05; 1K08AI146267-0) and a Cystic Fibrosis Foundation Harry Shwachman Award during the course of this study. This work was supported by the Ceragenin Research Fund granted to M.A.M. and J.S.C. by Bill Brown and Sharon Bonner-Brown.

## SUPPLEMENTAL MATERIALS

**Supplement Table S1.** Minimal inhibitory concentrations (MIC) of colistin, LL37, CSA13, CSA131, CSA44, CSA144 and ciprofloxacin against *E. coli, L. monocytogenes* and *Mycobacterium* spp.

**Supplement Figure S1**. Supplementary information related to CRISPRi screening.

Schematic of two examples of deep sequencing libraries from the strategy used to multiplex growth experiments using 50bp single end reads. Each library is barcoded using aTruSeq i7 index (orange), and additionally incorporates a 4bp barcode (teal) that is read out by the TruSeq Read 1 primer. Each index (i7 and 4bp barcode) is introduced by the sequencing library PCR primers, enabling easy multiplexing. Library A incorporates a random offset at the start of Read 1 (NN) to ensure sequence diversity during cluster generation.

**Data Set S1.** Transcriptomic response of *E. coli* exposed to colistin, LL37, CSA13 and CSA131 as determined by RNAseq. This file includes raw and normalized read counts, fold changes, *P* and adjusted *P* values as well as lists of genes significantly up- or down-regulated.

**Data Set S2.** Annotation term enrichment analysis for significantly up- and down-regulated genes in *E. coli* exposed to colistin, LL37, CSA13 and CSA131. The analysis was performed with the DAVID Bioinformatics Resources using terms from UniProt keywords, COG ontology, GO and the KEGG pathway.

**Data Set S3**. Lists and annotation term enrichment analysis of genes significantly up- and down-regulated in *E. coli* exposed to all antibiotics, CAMPs and ceragenins. The analysis was performed with the DAVID Bioinformatics Resources using terms from UniProt keywords, COG ontology, GO and the KEGG pathway.

**Data Set S4**. Analysis of cis-acting elements associated with genes significantly up- and down-regulated in *E. coli* exposed to colistin, LL37, CSA13 and CSA131. This file includes lists of genes with a least one binding site for sigma factors or transcription factors as well as numbers of non-redundant binding sites for each conditions. Fold changes for genes of the CpxR, PurR, RcsA and PhoB regulons are also included.

**Data Set S5.** Proteomic response of *E. coli* exposed to colistin and CSA13. This file includes fold changes and adjusted *P* values (including imputed values) as well as lists of proteins significantly up- and down-regulated following exposure to colistin and CSA13.

**Data Set S6.** Annotation term enrichment analysis for proteins up- and down-regulated in *E. coli* exposed to colistin or CSA13. The analysis was performed with the DAVID Bioinformatics Resources using terms from UniProt keywords, COG ontology, GO and the KEGG pathway.

**Data Set S7.** Changes in abundance and statistical significance associated with each strains of the CRISPRi library following exposure to colistin, LL37, CSA13 and CSA131.

