## Supplementary material for "Ceragenins and antimicrobial peptides kill bacteria through distinct mechanisms": Table S1

### TABLE S1. Susceptibility of *E. coli*, *L. monocytogenes* and *Mycobacterium* spp. to antibiotics

| Specie | Strain | MIC[a] (µg/mL) | | | | | | |
|---|---|---|---|---|---|---|---|---|
| | | COL | LL37 | CSA13 | CSA131 | CSA44 | CSA144 | CIP |
| *E. coli* | MG1655 | 0.25-1 | 16-32 | 2-4 | 2-4 | 2-4 | 4-16 | 0.015 |
| *L. monocytogenes* | 10403S | 128- >128 | 32-64 | 1-2 | 1-2 | 2-4 | 4-8 | 1 |
| *M. avium* | $mc^2$2500 | >128 | >128 | 128- >128 | >128 | ND | ND | 0.5-2 |
| | $mc^2$2500D6 | >128 | >128 | 32 | 64 | ND | ND | 0.125-0.25 |
| *M. marinum* | M | >128 | >128 | 32-128 | 64-128 | ND | ND | 0.25-1 |
| *M. smegmatis* | $mc^2$155 | 64-128 | >128 | 0.125-0.25 | 0.5 | 16-32 | 64-128 | 0.25 |
| *M. tuberculosis* | Erdman | 128- >128 | >128 | 16 | 64 | ND | ND | 0.5 |

[a] COL, colistin; CIP, ciprofloxacin; ND, not determined.
