## Supplementary material for "Ceragenins and antimicrobial peptides kill bacteria through distinct mechanisms": Fig S1

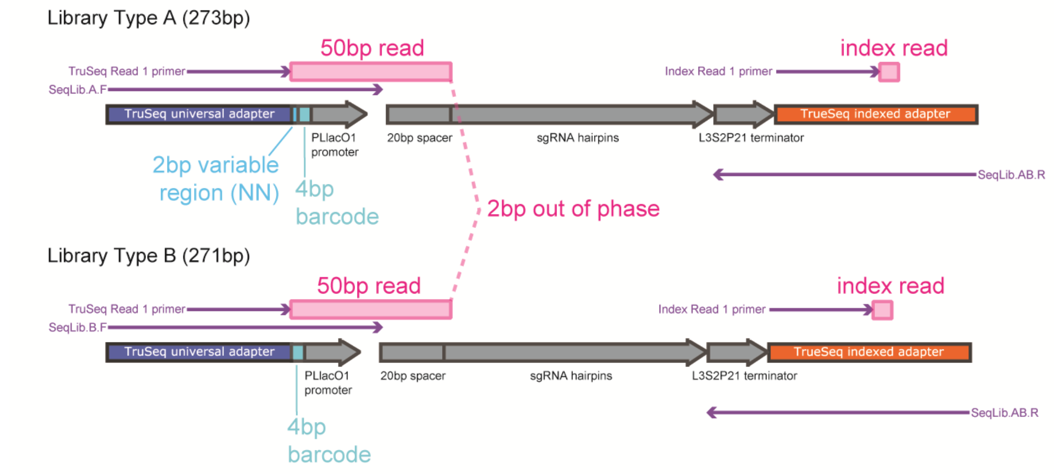


**FIGURE S1.** Supplementary information related to CRISPRi screening. Schematic of two examples of deep sequencing libraries from the strategy used to multiplex growth experiments using 50bp single end reads. Each library is barcoded using aTruSeq i7 index (orange), and additionally incorporates a 4bp barcode (teal) that is read out by the TruSeq Read 1 primer. Each index (i7 and 4bp barcode) is introduced by the sequencing library PCR primers, enabling easy multiplexing. Library A incorporates a random offset at the start of Read 1 (NN) to ensure sequence diversity during cluster generation.
